# (Pro)renin receptor inhibited ABCA1/G1 expression associated with low HDL-c and promoted atherosclerosis

**DOI:** 10.1101/2024.04.03.588014

**Authors:** Yubing Dong, Wei Zhang, Wanyu Liu, Cuiping Bai, Ying Zhang, Yinong Jiang, Wei Song

**Affiliations:** Department of Hypertension, The First Affiliated Hospital of Dalian Medical University, Dalian Liaoning 116011, China

**Keywords:** (Pro)renin receptor, ATP-binding cassette transporter A1, ATP-binding cassette transporter G1, atherosclerosis

## Abstract

**Background:** (Pro)renin receptors [(P)RR] binding both renin and its inactive proenzyme for prorenin induce increasing the efficiency of angiotensin I conversion and the production of angiotensin II (Ang II). Recently, (P)RR was reported to played a pathological role in diabetic renal disease independent of Ang II. Ang II is involved in lipid metabolism and atherosclerosis, but the (P)RR participation in atherosclerosis remains unclear. Thus, the objective of this study was to examine the relative importance of Ang II-independent actions of (P)RR and its mechanism in the development of atherosclerosis.

**Methods:** Clinical demographic characteristics and laboratory profile data of 1189 essential hypertensive were collected. Patients were divided into four groups based on the quartiles of plasma renin activity (PRA). The plasma lipoprotein levels of four groups were compared. Furthermore, we examined serum and liver lipoprotein levels, as well as ATP-binding cassette transporter A1 (ABCA1) and ATP-binding cassette transporter G1 (ABCG1) expression in ApoE^-/-^ mice treated with prorenin. ABCA1 and ABCG1 expression and their function in mediating cholesterol efflux were evaluated in THP-1 cells by prorenin treatment. Agonists of the peroxisome proliferator-activated receptor-γ (PPAR-γ) and liver X receptor α (LXRα) were used to investigate the mechanism involved in vitro.

**Results:** Plasma TC, TG, LDL-c, and apoB were significantly increased, and HDL-c and apoA were dramatically decreased with a trend of higher plasma renin activity (PRA) ( all *P* < 0.05) in hypertensive patients. Prorenin accelerated atherosclerosis by inhibiting mRNA and protein expression of ABCA1/ABCG1 in the liver of ApoE^-^/^-^ mice. Prorenin binding to (P)RR inhibited the functions of ABCA1- and ABCG1-mediated cholesterol efflux. Furthermore, Prorenin reduced ABCA1/G1 expression of THP-1 cells through PPARγ/LXRα pathway.

**Conclusions:** (P)RR activated by prorenin inhibited the expression of ABCA1/ABCG1 through the PPARγ/LXRα pathway and the functions of ABCA1- and ABCG1-mediated cholesterol efflux, resulting in dyslipidemia and eventually atherosclerosis.

**Highlights:** With RPA increases in hypertension, plasma TC, TG, LDL-c, and apoB were higher, while HDL-c and apoA were lower.

Prorenin binding to (P)RR accelerated atherosclerosis by inhibiting ABCA1/G1 expression and function.

Prorenin reduced ABCA1/G1 expression of THP-1 cells through PPARγ/LXRα pathway.

## INTRODUCTION

Atherosclerosis is the primary inducer of myocardial infarction, ischaemic cardiomyopathy, strokes, and peripheral arterial disease, and has become prevalent in more populous developing countries nowadays^1^. It is characterized by an elevation in plasma low-density lipoprotein (LDL) cholesterol, triglyceride (TG), and a decrease in high-density lipoprotein (HDL)^1^. The accumulation of cholesterol in intravascular macrophages is a crucial factor for the progression of atherosclerosis. HDL involves a protective mechanism against atherosclerosis, called reverse cholesterol transport (RCT), through the transfer of excess cholesterol from macrophages and peripheral tissues to the liver, where the cholesterol is eventually metabolised and excreted via the bile and faeces^2^. Consequently, stimulation of HDL-c levels may provide a therapeutic strategy for atherosclerosis.

The ATP-binding cassette transporter A1 (ABCA1) and the ATP-binding cassette transporter G1 (ABCG1) are members of the ABC transmembrane transporter family. ABCA1 and ABCG1-mediated cholesterol efflux is the initiation step of RCT. ABCA1 mediates cholesterol efflux from macrophages to lipid-free apolipoprotein A-I (apoA-I) was a limiting step in promoting nascent HDL generation^3^. ABCA1 gene mutation causes autosomal recessive genetic disorder Tangier disease, in which patients show extremely low plasma levels of HDL-c, thus leading to a high risk for atherosclerosis^4^. ABCG1 also plays a role in maintaining cellular cholesterol homeostasis by facilitating the efflux of intracellular cholesterol to nascent HDL^2,5^. A study showed that ABCG1 knockout mice on a high-cholesterol diet had a markedly decreased plasma HDL-c level^6^. Additionally peroxisome proliferator-activated receptor (PPAR)γ agonists can activate liver X receptor (LXR)α, resulting in increased expression of ABCA1 and ABCG1^7^.

As a new member of the renin-angiotensin system (RAS) family, (P)RR is a co-receptor for prorenin and renin. Prorenin is converted to active mature renin upon binding to (P)RR. When mature renin binds with the (P)RR, it facilitates the production of angiotensin II (Ang II) by promoting the effective conversion of angiotensinogen to angiotensinogen I^8,9^. Ang II is closely associated with the pathogenesis of atherosclerosis. Previous studies found that Ang II binding into Angiotensin I type receptor (AT1R) can activate the activation of cytokines such as TNF-α and IL-6, chemokines like MCP-1, and adhesion molecules including P-selectin, ICAM-1, and VCAM-1. These molecules play a crucial role in promoting the migration of mononuclear leukocytes and causing vascular inflammation.

Angiotensin also can facilitate the oxidation of LDL-C, leading to the creation of foam cells^10^. Therefore, the traditional view suggests that (P)RR is indirectly involved in the pathological process of atherosclerosis through the production of Ang II. However, recent studies confirmed that rats overexpressed the (P)RR gene showed proteinuria, glomerulosclerosis, and increased levels of cyclooxygenase-2. These effects are not dependent on Ang II production^11,12^. It remains unclear if the (P)RR can contribute to the development of atherosclerosis independently of Ang II production. The present study was designed to explore the association between (P)RR independent of Ang II and HDL-C in the development of atherosclerosis.

## MATERIALS AND METHODS

### Study subjects and groups

A total of 1189 patients with essential hypertension who were admitted to the First Affiliated Hospital of Dalian Medical University between January 2011 and December 2021 were enrolled. Ethical approval was obtained from the Human Ethics Committee of the First Affiliated Hospital of Dalian Medical University, and all patients enrolled in the study signed an informed consent form. The inclusion criteria were 1) a diagnosis of essential hypertension was satisfied^13^; 2) age between 18 and 80 years; 3) all patients were completely drug-eliminated (angiotensin-converting enzyme inhibitors, angiotensin receptor antagonists, β-blockers and dihydropyridine calcium channel blockers were discontinued for at least 2 weeks, diuretics for at least 4 weeks and aldosterone receptor antagonists for at least 6 weeks (α-blockers or non-dihydropyridine calcium channel blockers may be used to control blood pressure during the withdrawal period). The exclusion criteria were 1) secondary hypertension (primary aldosteronism, renal parenchymal hypertension, renal artery stenosis, Cushing’s syndrome, pheochromocytoma, etc.), white coat hypertension, pseudo hypertension; 2) severe hepatic insufficiency (transaminases more than 3 times the upper limit of normal) and severe renal insufficiency (eGFR < 60 ml/min/1.73 m²); 3) women who have recently taken contraceptives and pregnant women; 4) patients who have recently taken ezetimibe and fenofibrate.

We determined Ang I concentrations in ambulatory plasma by radioimmunoassay and used Ang I levels as a proxy for plasma renin activity (PRA) levels^14^. The enrolled patients were divided according to the quartile method according to PRA levels into groups Q1 (PRA ≤ 0.39 ng/ml/h, n = 300), Q2 (0.39 < PRA ≤ 0.90 ng/ml/h, n = 301), Q3 (0.90 < PRA ≤ 2.07 ng/ml/h, n = 290), and Q4 (PRA > 2.07 ng/ml/ h, n = 298).

### Clinical Assessment

Clinical data were collected including age, duration of hypertension, gender, family history of hypertension, body mass index (BMI), history of smoking, history of alcohol consumption, history of coronary heart disease, history of stroke, and history of diabetes mellitus. Blood was taken from enrolled patients during their hospital stay and plasma levels of total cholesterol (TC), triglycerides (TG), HDL-c, LDL-c, apoA, and apolipoprotein B (apoB) were measured using Hitachi 7170 automatic biochemical analyzer (WEI RIKANG Bioengineering Co. Ltd., China). BMI = height (kg)/weight^2^ (cm^2^).

### Animals and Experimental Design

Six-week-old male ApoE^-/-^ C57BL/6 mice (Changsheng Biological, Shenyang, China) were maintained on a 12-hour light-dark cycle at room temperature under pathogen-free conditions and free access to normal laboratory diet and water. Eight-week-old C57BL/6 mice were fed an atherogenic diet (1% cholesterol and 20% fat) for eight weeks. At the same time, the mice were administered losartan (20 mg/kg/d) by gavage; and given prorenin (0.1 mg/kg) in the model group or equal doses of saline in the control group for eight weeks by implantable capsule osmolarity micropump (XMJ Scientific, Beijing, China). All studies followed the guidelines for the management and use of laboratory animals established by the National Institutes of Health Graduate School (NIH Publication No. 85-23, revised 1996) and approved by the Laboratory Animal Management Committee of Dalian Medical University.

### Biochemical Analyses

Mice fasted for 12 hours and flowed by retroorbital blood withdrawal. Plasma and liver TC, TG, LDL-c, and HDL-c were measured using a commercial kit (Jiancheng Biological, Nanjing, China). The sPRR, CRP, and prorenin/renin concentrations in mice serum were detected according to the manufacturer’s instructions (Meibiao, Jiangsu, China).

### Assessment of Atherosclerotic Plaques

Mice were euthanized, and the thoracic cavity was opened. The heart and aorta were flushed with saline. Hearts were fixed in 4% paraformaldehyde for 24 hours and more, followed by gradient dehydration in sucrose solution. Hearts were embedded in OCT (Sakura Finetek Co. Ltd., Japan), followed by rapid freezing, set to a section thickness of 7 μm, and serially sectioned at the root of the aorta. Oil-red O (Sigma-Aldrich, St. Louis, MO) staining was used to visualise the atherosclerotic load in the aortic root.

### Quantitative Real-Time PCR

Trizol (Invitrogen, California, USA) was used to extract total RNA, followed by reverse transcription using a kit (YEASEN, Shanghai, China). SYBR Green PCR reagent (Takara Co. Ltd., Japan) was chosen for real-time fluorescence quantification. The following real-time PCR primer sequences were used: ABCA1 forward,

5′-GGAGCCTTTGTGGAACTCTTCC-3′ and reverse

5′-CGCTCTCTTCAGCCACTTTGAG-3′; ABCG1 forward,

5′-GACACCGATGTGAACCCGTTTC-3′ and reverse

5′-GCATGATGCTGAGGAAGGTCCT-3′; GAPDH forward, 5′-

CATCACTGCCACCCAGAAGACTG-3′ and reverse 5′-ATGCCAGTGAGCTTCCCGTTCAG-3′. The 2^-ΔΔCt^ method was elected to analyse gene expression and was normalized to GAPDH as an internal control.

### Immunoblotting

Protein lysates were separated by electrophoresis in 8-12% sodium dodecyl sulfate-polyacrylamide gel electrophoresis (SDS-PAGE) gels. The following antibodies were selected: rabbit anti-renin receptor, rabbit anti-ABCA1, rabbit anti-ABCG1 (Abcam, Cambridge, UK), rabbit anti-PPARγ (Ebtek, Wuhan, China), rabbit anti-LXRα (Ebtek, Wuhan, China), rabbit anti-GAPDH (HUABIO, Hangzhou, China). After incubation with secondary antibodies, protein blots were detected using ECL luminescent solution (Bio-Rad, CA, USA). Blots were quantified using ImageJ (ImageJ Software, NIH, Bethesda, USA) and normalised using GAPDH as an internal control.

### THP-1 culture and treatment

Human THP-1 macrophages (KGI, Jiangsu, China) were cultured in RPMI-1640 containing 10% fetal bovine serum at 37 °C in a humidified atmosphere of 5% CO_2_. Cells were differentiated into macrophages after incubation with 100 ng/ml phorbol myristate acetate for 72 h.

### Construction of (P)RR siRNA transfection and overexpression models

The specific siRNA against (P)RR (sense, 5′-GGGAACGAGUUUUAGUAUAUTT-3′; antisense, 5′-AUAUAUACUAAACUCGUUCCCTT-3′) and the negative control siRNA (sense, 5′-UUCUCCGAACGUGUCACGUTT -3′; antisense, 5′-ACGUGACACGUUCGGAGAATT-3′) were synthesized (Jima, Suzhou, China). The recombinant viral plasmid encoding the lentiviral particles and the negative control were purchased from Jima (Suzhou, China). The lentivirus was selected to insert a full-length 1053 bp human-derived (P)RR gene between cloning sites NotⅠ/BamHⅠ. Immunoblotting was conducted to detect transfection and overexpression efficiencies.

### Cellular cholesterol efflux assay

THP-1 macrophages were cultured with or without 2.5 x 10^-8^ mol/l prorenin for 72 h. After incubation, cells were incubated with 5 μmol/l 22-NBD-cholesterol mixed with medium for 4 h. After washing with PBS, cells were then incubated with 30ug/ml of apoA1 or 50 μg/ml of HDL induction effluent for 4 h. Induction effluent and cell lysate were collected, respectively. The fluorescence intensity (FI) of the medium and lysate was measured using an enzymatic calibrator (Perkin Elmer, Massachusetts, USA) at 469 nm excitation wavelength and 537 nm emission wavelength. Finally, the percentage efflux was calculated using the following equation: cholesterol efflux rate = (induced effluent FI/ (induced effluent FI + cell lysate FI)) × 100%.

### ABCA1 mRNA stability assay

THP-1 macrophages with or without overexpression of (P)RR were intervened with prorenin at a concentration of 2.5 x 10^-8^ mol/l for 72 h, then actinomycin D was added, and cellular RNA was collected at 0, 2, 4, 6, and 8 h to detect ABCA1 mRNA levels.

### Statistical Analysis

Clinical data were obtained using SPSS software version 26.0 (SPSS Inc., Chicago, IL, United States). If the variables were continuous and normally distributed, ANOVA was performed, and the results were presented as mean ± standard deviation. If not normally distributed, the Kruskal-Wallis H test was used, and the results were described as median and interquartile. The chi-square test was applied for qualitative data, and the results were expressed as percentages. Correlations between variables were studied using Pearson or Spearman correlation analysis. Multiple linear stepwise regression was used to study the relationship between TC, TG, HDL-c, LDL-c, and PRA. Student’s t-test was performed on experimental data using Prism9 software (GraphPad Software, San Diego, CA, USA). Data were presented as mean ± standard deviation, and P < 0.05 was considered statistically significant.

## RESULTS

### Demographic characteristics and the relationship of lipid profile to PRA

There was no significant difference between the four groups regarding the course of hypertension, family history of hypertension, history of smoking, alcohol consumption, cerebral infarction, and diabetes mellitus (each *P* > 0.05). The four groups exhibited differences in age, sex, body mass index, and the course of coronary heart disease (each *P* < 0.05) (Table S1). TC, TG, LDL-c, and apoB were significantly increased with a trend of higher plasma renin activity (PRA) (*P* = 0.011, *P* < 0.001, *P* = 0.001, and *P* = 0.003). However, HDL-c and apoA were dramatically decreased with a trend of higher PRA (*P* = 0.014 and *P* = 0.028) (Table 1). The association between the TC, LDL-c, HDL-c, and PRA levels remained after adjustment for age, sex, BMI, and coronary heart disease.

**Table 1.** Comparison of lipid profiles in the four groups.

|  | Q1 | Q2 | Q3 | Q4 | Total | P |
| --- | --- | --- | --- | --- | --- | --- |
|  | (n=300) | (n=301) | (n=290) | (n=298) | (n=1189) | Value |
| TC | 4.67±0.90 | 4.80±0.90 | 4.87±0.89 | 4.90±0.88 | 4.81±0.94 | 0.011 |
| (mmol/L) |  |  |  |  |  |  |
| TG | 1.26 | 1.46 | 1.52 | 1.57 | 1.44 | < 0.001 |
| (mmol/L) | (0.92,1.92 | (1.08,1.46 | (1.11,2.30 | (1.15,2.30 | (1.05,2.13 |  |
|  | ) | ) | ) | ) | ) |  |
| HDL-c | 1.13±0.25 | 1.10±0.23 | 1.10±0.20 | 1.07±0.20 | 1.10±0.25 | 0.014 |
| (mmol/L) |  |  |  |  |  |  |
| LDL-c | 2.62±0.64 | 2.74±0.66 | 2.81±0.63 | 2.81±0.62 | 2.74±0.64 | 0.001 |
| (mmol/L) |  |  |  |  |  |  |
| apoA | 1.05±0.21 | 1.02±0.19 | 1.02±0.19 | 1.00±0.17 | 1.02±0.19 | 0.028 |
| (mmol/L) |  |  |  |  |  |  |
| apoB | 0.88±0.27 | 0.92±0.20 | 0.93±0.19 | 0.95±0.21 | 0.92±0.22 | 0.003 |
| (mmol/L) |  |  |  |  |  |  |

TC, total cholesterol; TG, triglycerides; HDL-c, high-density lipoprotein cholesterol; LDL-c, low-density lipoprotein cholesterol; apoA, apolipoprotein A; apoB, apolipoprotein B. Data are presented as mean ± SD and median (interquartile range).

Multiple linear regression analysis revealed a positive relationship of PRA with TC, LDL-c (*P* = 0.031 and *P* = 0.028, respectively), and a negative relationship with HDL-c (*P* = 0.037), after adjustment for age, sex, BMI, smoking, and diabetes (Table 2).

**Table 2.**
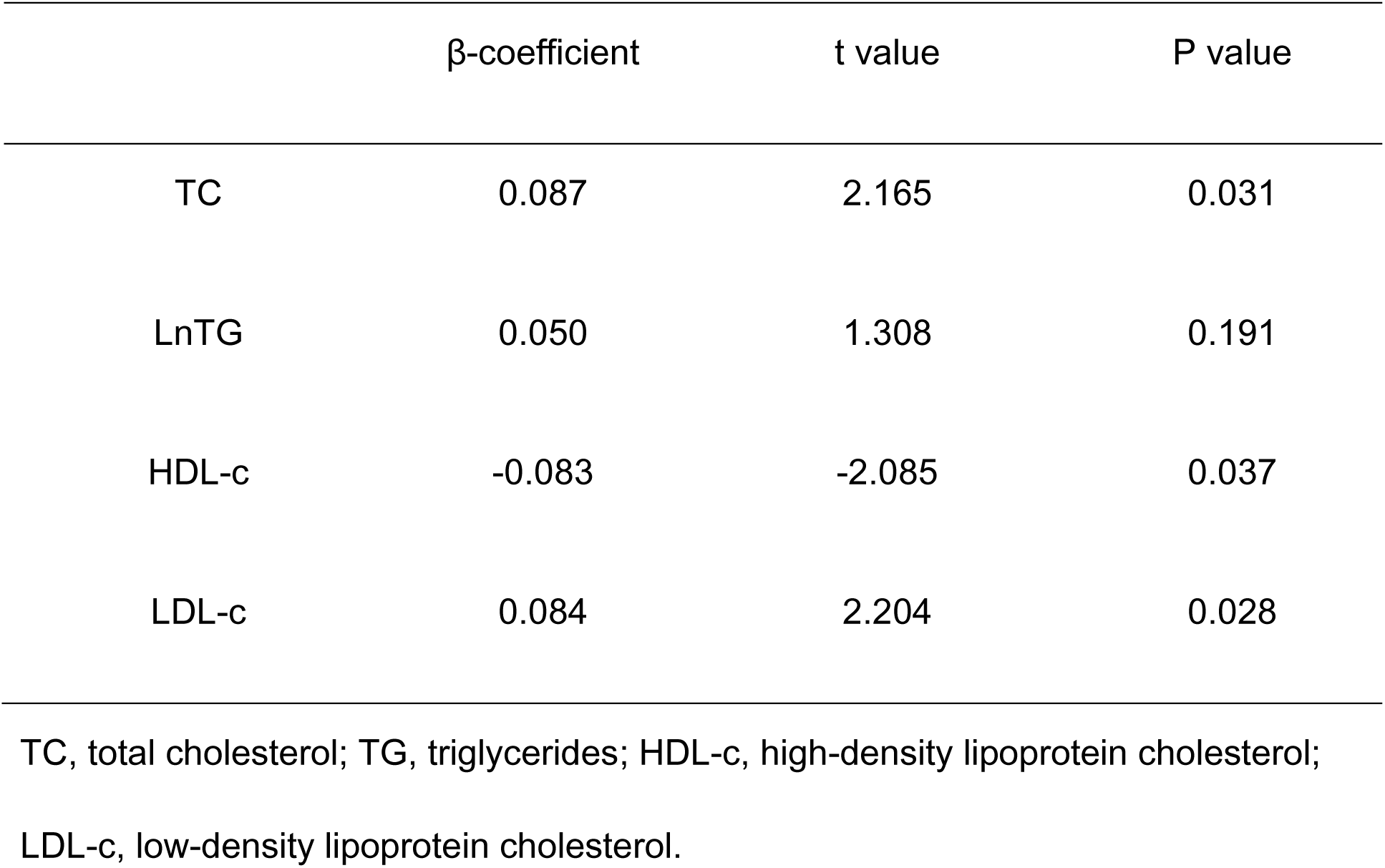
Multiple linear regression analysis testing the association of the lipid profiles.

| | β-coefficient | t value | P value |
| --- | --- | --- | --- |
| TC | 0.087 | 2.165 | 0.031 |
| LnTG | 0.050 | 1.308 | 0.191 |
| HDL-c | -0.083 | -2.085 | 0.037 |
| LDL-c | 0.084 | 2.204 | 0.028 |

TC, total cholesterol; TG, triglycerides; HDL-c, high-density lipoprotein cholesterol; LDL-c, low-density lipoprotein cholesterol.

### (P)RR accelerated AS by inhibiting mRNA and protein expression of ABCA1/ABCG1 in the liver of ApoE^-^/^-^ mice

To investigate the importance of (P)RR for atherosclerotic lesion development, **ApoE^-^/^-^** mice feeding atherogenic diet treated with prorenin and losartan were used as the prorenin group, whereas saline and losartan were used as control. Plasma levels of prorenin/renin, sPRR, and CRP were significantly increased in the prorenin group (*P* = 0.017, *P* = 0.004, and *P* = 0.004, respectively) (Figure S1A-C). In addition, body weight and systolic blood pressure were not statistically different between the two groups (*P* > 0.05) (Figure S1D-E).

Prorenin-treated mice had an increased aortic root lesion area (*P* = 0.001) (Figure 1A). The plasma levels of TG and LDL-c were significantly increased in the prorenin group compared to the control group (both *P* = 0.001). Meanwhile, the plasma levels of HDL-c were markedly lower in the prorenin group (*P* = 0.018) (Figure 1B). The liver lipid profiles demonstrated that TC and LDL-c levels were significantly increased in the prorenin group (both *P* < 0.001). However, HDL-c levels were dramatically reduced in the prorenin group (*P* = 0.024) (Figure 1C). The above results indicated that plasma and liver lipid profile changes affected by prorenin were consistent. We also found that liver ABCA1 and ABCG1 mRNA expression were significantly reduced in the prorenin group (*P* = 0.023 and *P* < 0.001 respectively) (Figure 1D and 1E). Meanwhile, liver ABCA1 and ABCG1 protein expression were also significantly suppressed in the prorenin group (*P* = 0.023 and *P* = 0.007 respectively) (Figure 1F and 1G).

**Figure 1.**
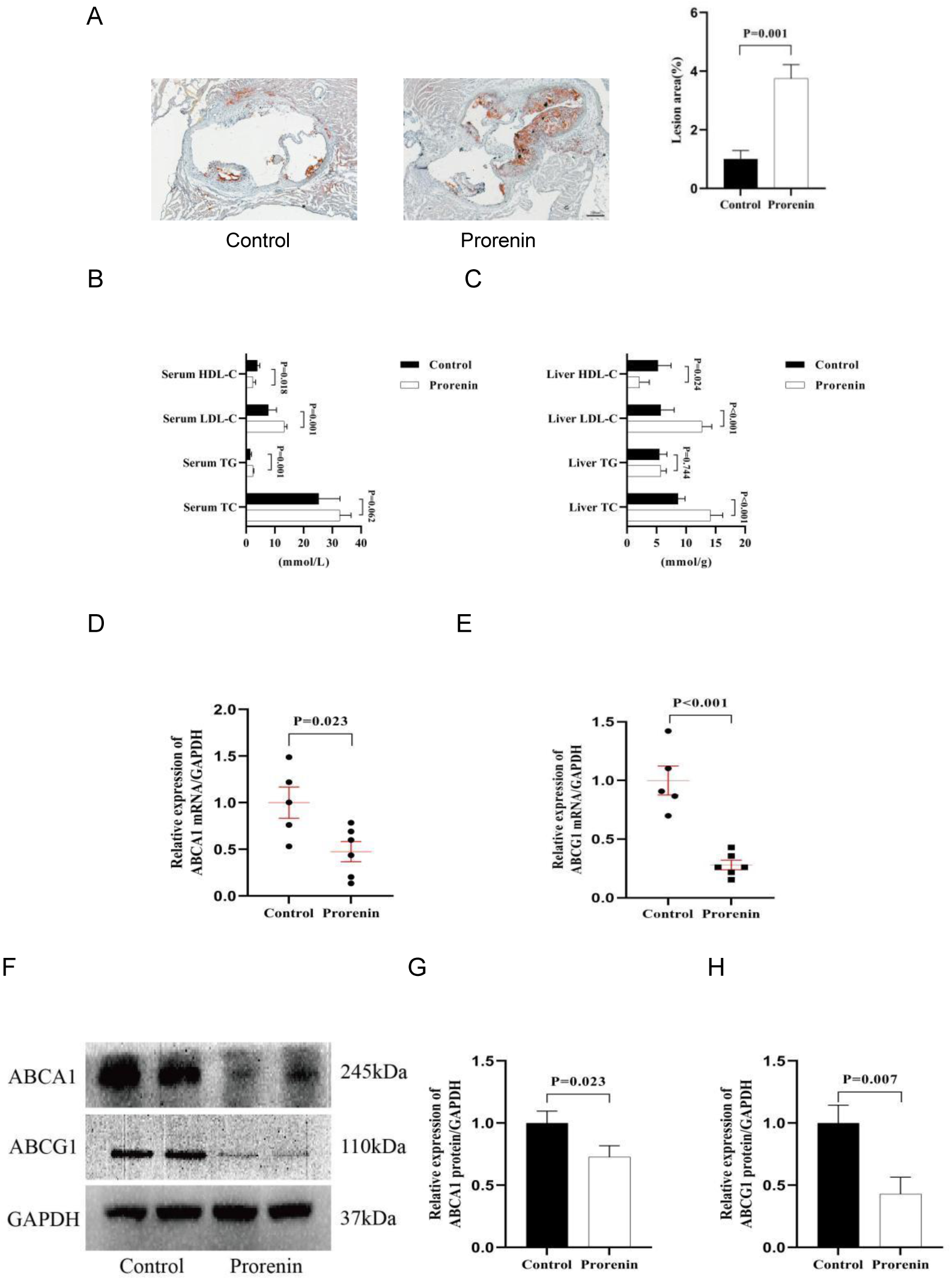
(P) RR accelerated aortic atherosclerosis in apoE^-^/^-^ mice. (A) Representative images of Oil Red O staining in the frozen section of the left ventricular outflow tract (scale bar, 200 μm) and quantification of lesion area. (B, C) total triglyceride (TG), total cholesterol (TC), low-density lipoprotein cholesterol (LDL-c) and high-density lipoprotein cholesterol (HDL-c) concentrations of serum and livers were measured (n=5-6). (D, E) ABCA1/ABCG1 mRNA levels in the livers of apoE^-^/^-^ mice (N=3). (F-H) Protein levels and quantification of ABCA1 and ABCG1 (n=3). Values are mean ± SD.

### (P)RR inhibited ABCA1/ABCG1 expression and ABCA1/ABCG1-mediated cholesterol efflux in THP-1 cells

The expression of ABCA1 and ABCG1 mRNA was significantly inhibited in THP-1 cells incubated with prorenin (*P* = 0.001 and *P* < 0.001) (Figure 2A). The protein levels of ABCA1 and ABCG1 were significantly reduced in THP-1 cells by prorenin treatment (*P* = 0.037 and *P* = 0.049) (Figure 2B, C lane 1 and lane 2). (P)RR expression was significantly suppressed following (P)RR siRNA transfection (Figure S3A). The protein expression of ABCA1 and ABCG1 was reversed by the treatment of (P)RR siRNA (*P* = 0.006 and *P* = 0.001) (Figure 2B, C lane 2 and lane 3). Additionally, when (P)RR expression was markedly upregulated after LV-(P)RR transfection (Figure S3B), ABCA1 and ABCG1 were further inhibited in comparison to the prorenin group (*P* = 0.021 and *P* = 0.054) (Figure 2B, C lane 2 and lane 4). ABCA1-mediated cholesterol efflux to ApoA1 and ABCG1-mediated cholesterol efflux to HDL were markedly decreased by prorenin intervention compared to the control group (*P* = 0.003 and *P* = 0.001). When cells were transfected with (P)RR siRNA, the inhibitory effect of (P)RR on ABCA1 and ABCG1-mediated cholesterol efflux rate were reversed (both *P* < 0.001). However, ABCA1 and ABCG1-mediated cholesterol efflux rates were further inhibited after LV-(P)RR transfection (*P* = 0.013 and *P* = 0.001) (Figure 2D). These results indicated that overactivation of (P)RR significantly inhibited the expression of ABCA1 and ABCG1 in THP-1 macrophages and the function of ABCA1- and ABCG1-mediated cholesterol efflux to apoA1 and HDL.

**Figure2.**
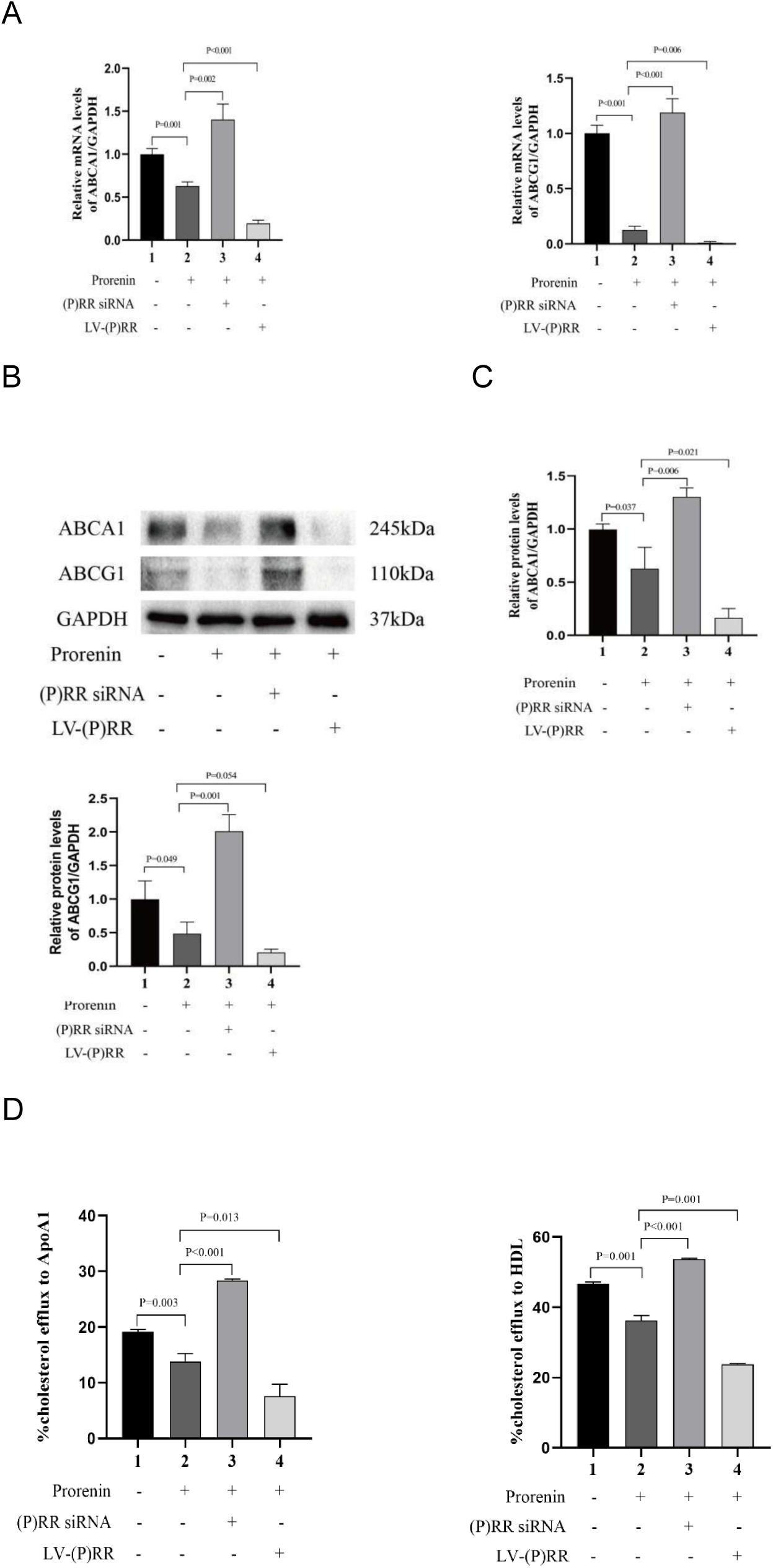
(P)RR inhibited the expression and function of ABCA1/ABCG1 in THP-1 macrophages. Cells were transformed with or without PRR siRNA or LV-(P)RR and pretreated with or without 2 × 10^-8^ mol/L prorenin for 72 h. (A) Quantification of mRNA of ABCA1 and ABCG1 in THP-1 cells. (B) Representative immunoblots of ABCA1/ABCG1 levels in THP-1 cells. (C) Quantification of protein expression of ABCA1 and ABCG1 in THP-1 cells. (D)The induced efflux and cell lysates were collected to measure the fluorescence intensity and calculate the cholesterol efflux rate (THP-1 macrophages were incubated with 5μmol/L 22-NBD-cholesterol for 4 h and ApoA1 (30 ug/mL) or HDL (50 μg/mL) was used to induce outflow as lipid acceptors for 4 h). N=3. Data were expressed as mean ± SD.

### (P)RR reduced the stability of ABCA1 mRNA through PPARγ/LXRα pathway

Prorenin stimulation of THP-1 macrophages decreased PPARγ and LXRα expression levels (*P* = 0.018 and *P* < 0.001), and also inhibited ABCA1 and ABCG1 expression (*P* < 0.001 and *P* = 0.001) (Figure 3A-E lane 1 and lane 2). PPARγ and LXRα protein expression were reversed by LXRα agonist (lane 3) or PPARγ agonist (lane 4) compared to prorenin (lane 2) (*P* < 0.05)(Figure A-E). ABCA1 and ABCG1 protein expression were similarly reversed by LXRα agonist (lane 3) or PPARγ agonist (lane 4) compared to prorenin (lane 2) (*P* < 0.05) (Figure 3A, D, E). In addition, the combination of LXRα and PPARγ agonists (lane5) further reversed PPARγ, LXRα, ABCA1, and ABCG1 protein expression compared to used alone (lane 3-4) (*P* < 0.05) (Figure 3A-E). These results demonstrated that (P)RR activated by prorenin could inhibit ABCA1 and ABCG1 expression through the PPARγ/LXRα pathway. We also found that ABCA1 mRNA expression in THP-1 macrophages after 72 h of prorenin intervention decreased progressively with increasing time of addition of actinomycin D and was less in all than in the control group, suggesting that (P)RR decreased the mRNA stability of ABCA1 (Figure 3F).

**Figure3.**
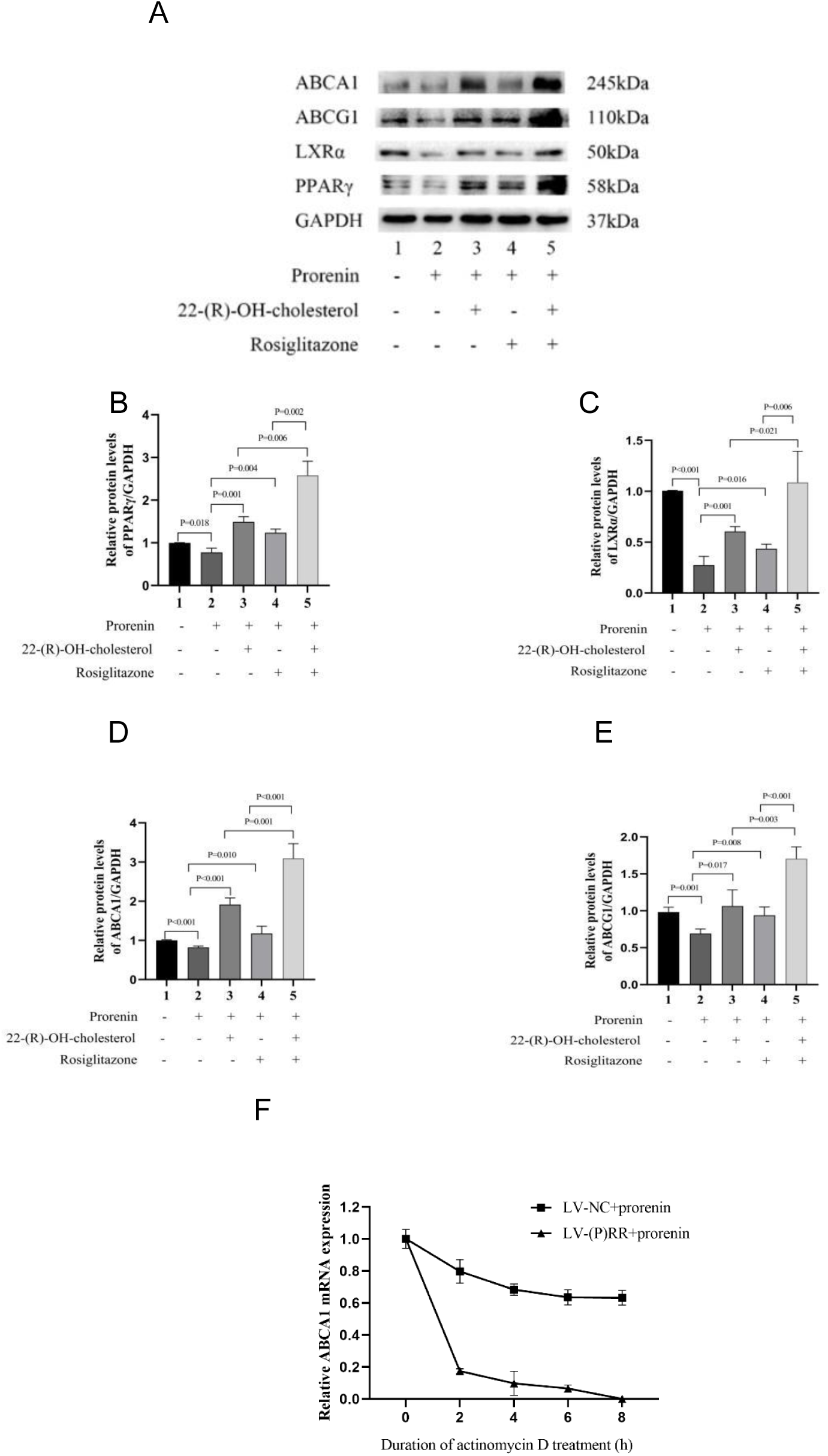
(P)RR decreased ABCA1 mRNA stability through PPARγ/LXRα pathway. (A) The protein expression of PPARγ, LXRα, ABCA1 and ABCG1 (prorenin intervention in macrophages was followed by incubation with 22-(R)-OH-cholesterol (1 μmol/L) or rosiglitazone (100 nmol/L) for 24 h). (B-E) Quantification of protein expression of PPARγ, LXRα, ABCA1 and ABCG1. (F) Detection of ABCA1 mRNA expression at each time period (prorenin intervention on macrophages for 72 h followed by treatment of cells with actinomycin D (final concentration 5 μg/ml). N=3. Data were expressed as mean ± SD.

## Discussion

(P)RR has been implicated as the receptor for renin/prorenin and acts by activating the renin-angiotensin system. It is becoming increasingly clear that Ang II may participate in the metabolism of lipids and the development of atherosclerosis through several mechanisms. The role of (P)RR in lipoprotein metabolism independent of AngII has not been elucidated. A previous study demonstrated that (P)RR regulates LDL metabolism independently of the RAS by controlling sortilin-1 and LDL receptor levels at the cellular level^15^. In this study, we found that PRA showed a positive relationship with TC and LDL-c and a negative relationship with HDL-c, suggesting PRA may involved in dyslipidemia. These results were consistent with a previous study of hypertension patients from 261 young men, which found a significant positive correlation between LDL-C and PRA^16^. A study was conducted on mild hypertension patients, which included 58 patients with high PRA (PRA>0.65ng/ml/h) and 39 patients with low PRA (PRA≤0.65ng/ml/h)^17^. This study found that the high PRA group had significantly higher TC and LDL-C levels than the low PRA group, partially consistent with our results. Additionally, we also found that the high PRA group had significantly lower HDL-C levels than the low PRA group.

In this study, aortic atherosclerotic plaques significantly increase in mice subcutaneously pumped with prorenin, suggesting that (P)RR activated by prorenin can exacerbate AS. It has been consistently demonstrated that Ang II binding to AT1R accelerates the development of atherosclerosis^18–20^. The pharmacological AT1R inhibitor, losartan, reduced experimental atherosclerosis and blood pressure in a similar dose-dependent manner^21^. In this study, we inhibited the effect of Ang II by losartan on two groups of mice. The effect of Ang II was abolished by losartan, given that the blood pressure of mice in both groups did not increase. It suggested that the aggravation of atherosclerosis in the prorenin group did not result from the interaction between Ang II and ATIR. Nguyen et al. proposed that prorenin receptors binding with prorenin can activate the ERK1/ERK2 pathway independently of the RAS activation^22^. In this study, plasma levels of prorenin/renin and s(P)RR were significantly increased in the prorenin group, indicating that the increased level of prorenin activated the expression of its receptor (P)RR. Thus, accelerated AS by (P)RR is mediated through a RAS-independent mechanism.

It is well known that the increase of LDL-C and the decrease of HDL-C in plasma are the dyslipidemia characteristics of coronary artery disease. In this study, we found that the plasma levels of TG and LDL-c were increased in the prorenin group, and the plasma level of HDL-c was decreased. Lipid profiles of the liver are also changing in line with those in plasma, indicating that prorenin can change blood lipid profiles by activating (P)RR to accelerate AS development. Current treatments for atherosclerosis utilize statin lipid-lowering drugs as the first choice^23^. However, many patients discontinue statin due to side effects, mainly muscle-related effects^24^. It has recently been discovered that RCT could be an attractive therapeutic approach. In atherosclerotic plaques, ABCA1 and ABCG1 are key receptors for RCT. 70% of total cholesterol efflux from macrophage foam cells occurs via ABCA1 and ABCG1.

ABCA1 and ABCG1 knockout mice accumulate numerous macrophage foam cells in tissues such as the lung, liver, spleen, and thymus^25^. The ABCA1 gene mutation can cause Tangier disease, characterized by low plasma HDL-C levels and excessive cholesterol accumulation in the reticuloendothelial system, which increases the risk of cardiovascular disease^26^. The role of ABCG1 in the development of AS is still controversial. LDL receptor knockout mice selectively deficient in macrophage ABCG1 feeding a high-cholesterol diet aggravate atherosclerotic lesion development^27^. However, Tarling et al. found that ApoE and ABCG1 knockout mice showed more macrophage apoptosis and smaller atherosclerotic lesions^28^. In this study we found that activation of (P)RR by prorenin reduces HDL-C levels in plasma and the liver by inhibiting ABCA1 and ABCG1, thereby accelerating AS development. Macrophages are the primary cells overloaded with cholesterol involvement in AS. To eliminate the interference of Ang in vivo and understand the detailed mechanism more clearly, we used THP-1 macrophages cultured in vitro. Angiotensinogen is the only precursor of all angiotensin peptides^29^. Angiotensinogen is not present in macrophages cultured in vitro. Hence, Ang II also does not exist. We found that the mRNA and protein expressions of ABCA1 and ABCG1 in THP-1 macrophages decreased as the prorenin concentration increased. ABCA1 mediates the rate-controlling step in HDL formation by promoting the efflux of cholesterol and phospholipids to ApoA1. Furthermore, oxidized low-density lipoprotein is a key to AS development; ABCG1 mainly mediates the oxidized sterols derived from it to efflux into HDL rather than ABCA1. And, only sterols effluxed through the ABCG1 pathway are efficiently taken up by hepatocytes^30^. We found prorenin inhibited ABCA1-mediated cholesterol efflux to ApoA1 and ABCG1-mediated cholesterol efflux to HDL.

PPAR and LXR are nuclear receptors activated by fatty acids and cholesterol derivatives, respectively, and they control the expression of a series of genes involved in lipid metabolism and inflammation^31^. Studies have shown that the expression of ABCA1 and ABCG1 can be regulated by activating the PPARγ/LXRα pathway, affecting the efflux of cholesterol from macrophages^32,33^. This mechanism plays an essential role in the development of AS. In the mechanism of Ang II promoting atherosclerosis, Takata et al. reported that Ang II reduced the expression of ABCA1 in vascular macrophages of LDLR^-/-^ mice by inducing Fra2 bind to the E-box motif of ABCA1 promoter, and ABCA1 mRNA stability was not changed^34^. However, our results showed that (P)RR can reduce the mRNA stability of ABCA1. Therefore, (P)RR and Ang II promote atherosclerosis in different ways.

In summary, we provide new evidence that (P)RR represses ABCA1 and ABCG1 expression through the PPARγ/LXRα pathway and is at least achieved by reducing mRNA stability. We demonstrated for the first time that (P)RR can participate in atherosclerosis development by altering blood lipid levels independently of Ang II.

## Acknowledgments

None.

## Sources of Funding

This research was supported by the National Natural Science Foundation of China (Grant 82270475).

## Disclosures

None.

## Notes

### Competing Interest Statement

The authors have declared no competing interest.

